# Theoretical insights into rotary mechanism of MotAB in the bacterial flagellar motor

**DOI:** 10.1101/2024.03.25.586605

**Authors:** Shintaroh Kubo, Yasushi Okada, Shoji Takada

## Abstract

Many bacteria enable locomotion by rotating their flagellum. It has been suggested that this rotation is realized by the rotary motion of the stator unit, MotAB, which is driven by proton transfer across the membrane. Recent cryo-electron microscopy studies have revealed a 5:2 MotAB configuration, in which a MotB dimer is encircled by a ring-shaped MotA pentamer. While the structure implicates the rotary motion of the MotA wheel around the MotB axle, the molecular mechanisms of rotary motion and how they are coupled with proton transfer across the membrane remain elusive. In this study, we built a structure-based computational model for *Campylobacter jejuni* MotAB, conducted comprehensive protonation state-dependent molecular dynamics simulations, and revealed a plausible proton-transfer coupled rotation pathway. The model assumes rotation-dependent proton transfer, in which proton uptake from the periplasmic side to the conserved aspartic acid in MotB is followed by proton hopping to the MotA proton-carrying site, followed by proton export to the cytoplasm. We suggest that, by maintaining two of the proton-carrying sites of MotA in the deprotonated state, the MotA pentamer robustly rotates by ∼36° per proton transfer across the membrane. Our results provide a structure-based mechanistic model of the rotary motion of MotAB in bacterial flagellar motors and provide insights into various ion-driven rotary molecular motors.

**Significance Statement:** This study aims to elucidate the mechanism by which bacteria move by rotating their flagella. The driving force for flagellar rotation is predicted to be driven by protons passing through the transmembrane protein MotAB, but the actual rotation mechanism has not yet been elucidated. Using advanced computational modeling and molecular dynamics simulations, we have elucidated the detailed processes by which proton translocation achieves the rotation of the bacterial flagellar motor. This work not only sheds light on the fundamental mechanisms of bacterial motility but also provides a framework for understanding similar ion-driven rotation mechanisms in other biological systems, potentially paving the way for new bioinspired technologies.

## Introduction

Flagella are filamentous structures that are ubiquitously used to propel many unicellular organisms (1). In some eukaryotes such as *Tetrahymena thermophila* (2,3) and *Chlamydomonas reinhardtii* (4), the flagellum is bent by the ATP-dependent action of dynein to generate propulsion. In contrast, numerous gram-negative and -positive bacteria, such as *Campylobacter jejuni* (*Cj*)(5), *Clostridium sporogenes* (6), *Flavobacterium johnsoniae* (7), and *Serratia marcescens* (8), generate propulsion by rotating their flagella, driven by an ion concentration gradient across the membrane (9,10). This propulsion is closely related to bacterial motility and virulence (11).

Rotary bacterial flagellar motors are embedded in the membrane and consist of a stator and rotor (12–17). Of the two parts, the ring-shaped stator is responsible for ion transport that powers the motor (18–20). In many bacteria, the stator comprises an array of MotAB units. Recently, high-resolution cryo-electron microscopy (cryo-EM) studies revealed a “5:2 configuration” of gram-negative *Campylobacter jejuni* (*Cj*) MotAB, in which dimeric MotB’s are encircled by pentameric MotA’s (Fig. 1a-c) (5). This 5:2 architecture together with its tethering to the surroundings implicates the rotary motion of the MotA wheel around the central MotB axle, making the “wheels within wheels” model of flagellar motors plausible; namely, many rotary MotA wheels within the stator wheel transmit torque to the C-ring of the flagellar rotor (21,22). However, to date, the rotary motion of MotAB has not been directly observed. The rotary mechanism of MotAB, which is critical for understanding the rotary mechanisms of bacterial flagellar motors, has not yet been elucidated (23).

**Figure 1.**
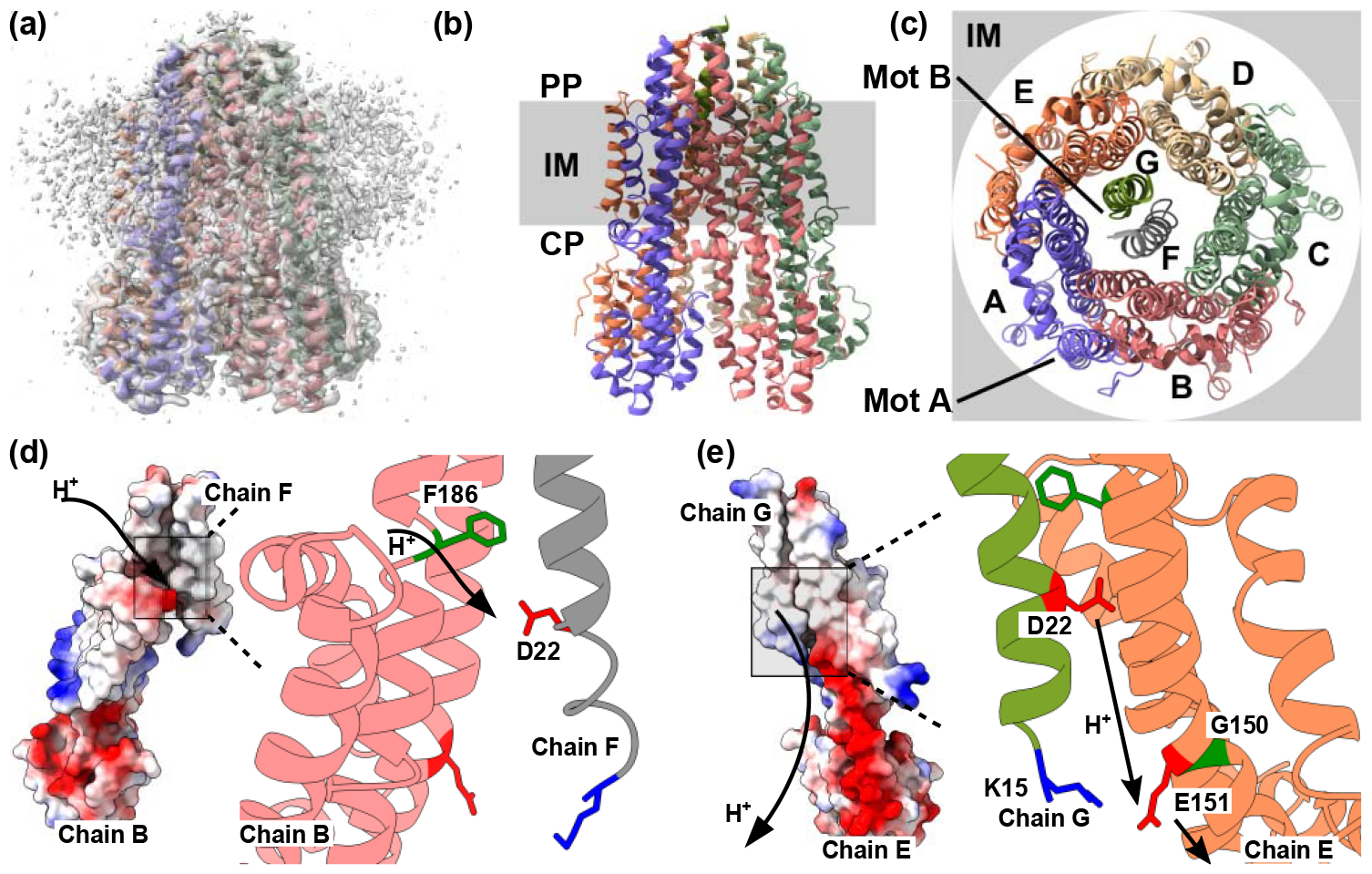
Structure of *Campylobacter jejuni* MotAB. (a) Cryo-EM structural model (PDB: 6YKP) depicts the overall structure of the MotAB. (b) Lateral view of the MotAB complex embedded in the inner membrane (IM). The top and bottom sides are periplasm (PP) and cytoplasm (CP), respectively. (c) Top view of the MotAB complex; MotA chains A to E are arranged counterclockwise. MotB chains F and G in the center serving as axle. (d) Proton uptake pathway from PP into IM. Left: Surface charge map of MotA (chain B) and MotB (chain F). Positive (negative) charges are in red (blue). Right: Cartoon representation of the corresponding region, with key residues drawn by sticks; F186 (green) in Chain B (salmon). D22 (red) in chain F (gray). (e) Possible proton transfer pathway from IM to CP side. Left: Surface charge map of MotA (chain E) and MotB (chain G). Right: Cartoon representation of the corresponding region, with key residues drawn by sticks; MotA (chain E salmon).E151; red. MotB (chain G green tea). D22; red and K15; blue. MotA (chain E).G150; green.

The central part of *Cj* MotAB structure was embedded in the inner membrane (Fig. 1, Fig. S1a). The proton motive force induces protons to flow from the periplasm to the cytoplasm (Fig. 1d,e), which is expected to drive pentameric MotA to rotate around the dimeric MotB. This is somewhat similar to other well-studied rotary motors, such as the Fo motor in F-type ATP synthase (24–26). Many structural (27–31) and computational (32–34) studies have investigated the driving Fo motor mechanisms. In the Fo motor, the c-ring (rotor) has highly conserved proton-carrying sites in the middle of the transmembrane helix, whereas the a-subunit (stator) has two half-channels of protons, each connected to either the cytoplasm or periplasm. A key feature is that the two half-channels of the stator (a-subunit) are not directly connected but can only be connected via the rotation of the proton-carrying sites in the rotor (c-ring). This can give a hint to the rotary “power-stroke” mechanism of MotAB (35,36).

The structural model of MotAB revealed a probable rotation-dependent proton uptake pathway. The side chain of MotA.F186 adopts multiple orientations depending on its interaction with MotB (Fig. S1b) (5). In particular, the orientation of MotA (chain B). F186 depicted in Fig. 1d and Fig. S1b is the only one that ensures an open proton uptake pathway from the periplasm to the middle of the inner membrane (the five subunits of MotA are designated as chains A through E, whereas the two MotB subunits are named chains F and G, as shown in Fig.1c). In other MotA subunits, F186 likely blocks proton uptake. Thus, the proton-uptake pathway depends on the rotary angle of the MotA wheel relative to that of MotB. The proton-carrying residue in the middle of the inner membrane in *Cj*MotB is believed to be D22 based on sequence conservation, mutagenesis experiments, and structural observations (Fig. 1d) (5,37,38). Notably, the proton enters the open gate of MotA(chain B).F186, which is near MotB(chain F).D22 in the cryo-EM structure. In the wild type (WT) cryo-EM structure, the side chain of MotB.D22 was oriented upward (toward the periplasm), whereas the side chain of D22N, introduced as a proxy for the protonated state, was oriented downward (toward the cytoplasm). Based on this, protonation of D22 has been hypothesized to cause the side chain to face the cytoplasm direction and release a proton toward the cytoplasm side. However, the subsequent proton release pathway towards the cytoplasm side and the coupling mechanism of proton transfer to rotary motion remains elusive.

In this study, we built a simple molecular dynamics (MD) simulation model for *Cj* MotAB to elucidate the rotary mechanism of ion-driven 5:2 symmetric motors. Based on the above argument for the cryo-EM structure, the model assumes that protons enter MotB(chain F).D22 from the periplasm-side, which was then transferred to the E151 moiety of MotA and released to the cytoplasm side (Fig. 1e). The E151 moiety includes the side chain of E151 and the backbone carbonyl group of G150, both of which can serve as proton (or hydronium ion)-carrying sites. Because of the 5:2 stoichiometry, there were seven protonatable sites in the simulation system: E151 in five MotA and D22 in two MotB. Thus, in principle, the model contained 2^7^ = 128 possible protonation states. Based on the investigation of cryo-EM structures, we simulated the transitions from one of these states to another to examine whether rotary motion occurred in the expected direction. Functional rotary motion was realized only when two of the five MotA.E151 states were deprotonated in the basal state. To the best of our knowledge, this is the first study to demonstrate the rotary motion of a MotAB using a concrete computational model. The results of this study will not only reveal the detailed rotation mechanism of MotAB but will also contribute to our understanding of the rotary mechanisms of ion-driven 5:2 symmetric motors.

## Results

### Modeling the proton-transfer coupled dynamics of MotAB

Our simulation was based on the *Cj*MotAB structure determined by cryo-EM (PDB ID: 6YKP) (5) and contained all the resolved regions of five MotA and two MotB molecules (Fig.1b). Based on the PDB structures, we annotated five MotA molecules as chains A to E in the counterclockwise direction from the periplasm side and two MotB molecules as chains F and G (Fig.1c). Reflecting that MotB is tethered in the flagellar stator, we restricted the MotB dimer to not rotate or translate and examined the motion of the MotA pentamer (see Methods for more details).

In our MD simulations for MotAB, we first modeled how the protons move: The proton uptake pathway from the periplasm side to D22 of MotB can, to some extent, be understood from the PDB structure. Namely, only MotB(chain B) had an unblocked side chain position of F186 for proton uptake (Fig.1d). MotB(chain B) is close to MotA(chain F) but not to chain G. Thus, it is reasonable to assume that the proton taken from the periplasm side reaches MotA(chain F).D22.

Given that, we focused on how protons are exported from MotB.D22 to the cytoplasm side. The surface charge of the *Cj*MotAB structure (PDB ID: 6YKP) is shown in Fig. 1de. One can clearly see a continuous negative-charged surface was observed from MotA.E151 to the cytoplasm side of MotA. Two potential proton-relaying sites were identified along the negative-charged path to the cytoplasm side. One is MotA.E151, which can transiently carry protons to form protonated E151 or attract hydronium ions. The negative charges at this site are moderately conserved (see Fig. S1 in (5)). Another potential site is the backbone carbonyl group of MotA.G150, which transiently captures hydronium ions. Notably, even in a helical structure, the carbonyl group of G150 cannot form a hydrogen bond with residue 154 because the latter is a highly conserved proline, P154. Thus, we assumed that a proton moved from MotB.D22 to either the side chain of MotA.E151 or the backbone carbonyl group of MotA.G150 (Fig.1e). Given the simple nature of our modeling, we did not distinguish between the two potential sites, and for simplicity, we designated these sites as E151. It is important to note that, from a functional point of view, this proton export is unlikely to occur in the same configuration as proton uptake, that is, from MotB(chain F).D22 to MotA(chain B).E151 in the configuration of the cryo-EM structure. If this were the case, the proton would have leaked without generating rotary motion. The most probable configuration for export is between MotB(chain G) and MotA(chain E) in the cryo-EM configuration because there is only one chain of MotB left, except for chain F, and because MotB(chain G) is the closest to MotA(chain E).

Inspecting the cryo-EM structure, we noticed that, in the vicinity of MotA(chain E).E151, there was a charged residue, MotB(chain G).K15 (Fig.1e). Thus, only when MotA(chain E).E151 was deprotonated, it was attracted to MotB(chain G).K15 via coulombic interactions. This attractive force was probably the strongest between MotA (chain E).E151 and MotB(chain G).K15 in the cryo-EM structure. It should be noted that this lysine is not well-conserved, but positively charged residues are found near this position in many species. This electrostatic attraction was expected to contribute to the enhancement of rotary motion.

Given these observations of the cryo-EM structure, we aimed to determine the proton transfer pathways that result in the rotary motion of the MotA pentamer around the MotB dimer. In this MD system, there were five MotA.E151 and two MotB.D22 protonatable residues, leading to a total of 2^7^ (128) possible protonation states. To reduce the number of likely states, we relied on the three propensities of the cryo-EM structure. 1) MotB(chain F).D22 is likely to be protonated because it is a proton uptake site. 2) Since proton then to jump to MotA(chain B).E151, we assumed that this site also tends to be protonated. 3) As previously discussed, MotA(chain E).E151 is close to MotB.(chain G).K15, indicating that MotA(chain E).E151 is most likely deprotonated.

With these propensities in mind, we explored the protonation state of MotA.E151. Motivated by this propensity, as the first trial of the simulation, we set the initial protonation state to be one in which only MotA(chain E).E151 was deprotonated, while all others were protonated (Fig. 2b, left cartoons). We denote this state as [MotA, MotB]=[(E), ()]; among five sites of MotA and two sites of MotB, deprotonated sites are indicated in the parentheses. From this reference, we gradually increased the number of deprotonated states in MotA one by one making a series of initial states, based on the above three propensities; a two deprotonated case ([(C, E), ()]), a three deprotonated case ([(A, C, E), ()]), a four deprotonated case ([(A, B, C, E), (G)]), where the propensity 3 is violated unavoidably (Fig. 2c-e,). For completeness, we also examined the zero-deprotonation state ([(), ()]) (Fig. 2a, where the propensity 2 is unavoidably violated).

**Fig. 2:**
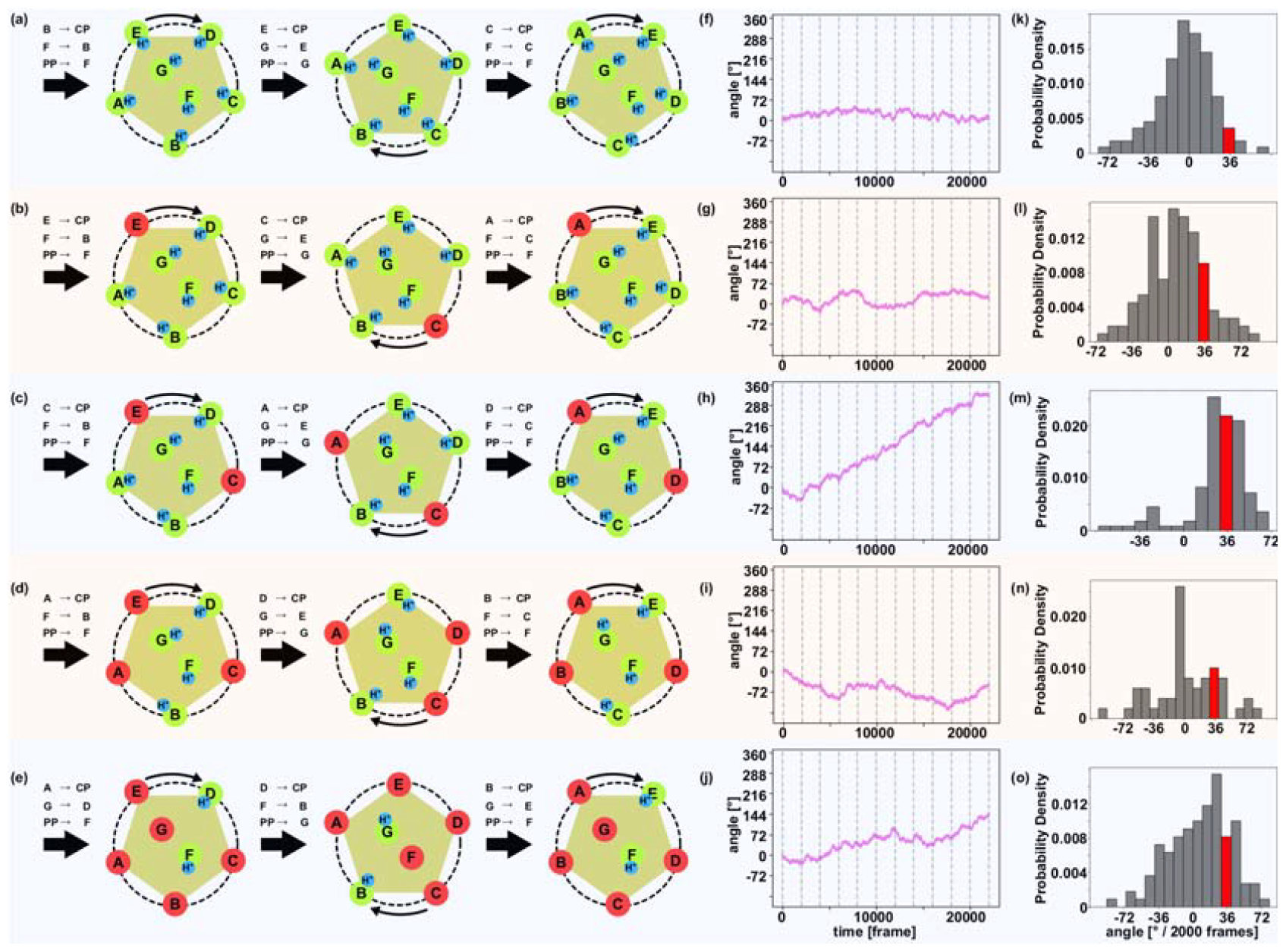
Rotary motions of MotAB with a varying number of deprotonated sites in MotA. (a-e) Diagrams showing different initial protonation states and their transitions during the simulation. Red circles represent deprotonated sites, and green circles indicate protonated sites. Protonated chains are adorned with a proton (blue). Initial states are (a) [MotA, MotB] = [(), ()], (b) [(E), ()], (c) [(C, E), ()], (d) [(A, C, E), (G)], and (e) [(A, B, C, E), (G)]. (f-j) Representative trajectories of rotary angles obtained from the simulations. The vertical dotted lines indicate updates of protonation states at every 2000 frames. (k-o) Histograms showing the probability density of the rotary angle change per update, calculating the rotary angle every 2000 frames. The red bin represents the 36°.

MD simulations were used to determine whether the MotA pentamer could rotate via successive cyclic updates of the protonation states (Fig. 2a-e). For MD simulations, we used a coarse-grained structure-based model, which has long been used to simulate large-scale protein motions with high efficiency and reasonable accuracy, using CafeMol software (39). In the model, each amino acid is represented by a single bead located at C_α_ position while pairwise interactions are calibrated based on atomic interaction at the native structure (see Methods for more details). Both the MotA pentamer and MotB dimer structures were separately biased towards their cryo-EM structures. While the position of the MotB dimer was restrained so that it does not rotate or translate, the MotA pentamer could move freely around the MotB dimer. Electrostatic and short-range repulsive interactions were applied between the MotA pentamer and MotB dimer.

In each simulation setup, we performed ten MD runs with different random numbers. In each trajectory, the protonated state was updated every 2000 × 10^4^ MD steps for a total of ten updates per trajectory (Fig. 2a-e shows a schematic diagram of the first three states and three updates). For convenience, 10^4^ MD steps were considered one frame and treated as the smallest unit of analysis.

### Rotary motion with the two deprotonated sites in the MotA pentamer

We performed comparative MD simulations of the five setups in Fig. 2a-e, starting from [MotA, MotB] = [(), ()] (Fig. 2a), [(E), ()] (Fig. 2b), [(C, E), ()] (Fig. 2c), [(A, C, E), ()] (Fig. 2d), and [(A, B, C, E), (G)] (Fig. 2e). Movies of representative trajectories in Fig.2c and Fig.2d are in Movie S1 and S2. The trajectories of the MotA pentamer rotation clearly show that robust rotary motions of the MotA pentamer appeared only in one case (representative trajectories in Fig. 2f-j, with all 10 trajectories in Fig. S2). In other words, only the setup starting from [MotA, MotB] = [(C, E), ()] led to a progressive rotation. For quantitative analysis, we plotted the average rotary angle change per protonation state update (every 2000 frames) (Fig. 2k-o). It was confirmed that only in the [(C, E), ()] setting, the most frequent rotation angle was approximately 36° per update of the protonation state.

We examined the rotary mechanism of MotAB when we started from [(C, E), ()], obtained a robust rotation (Fig. 2c, h, m) (we also performed the same simulations twice as long as the previous setup, confirming essentially the same results. See Fig. S3f, g, h). In this scheme, a single update of the protonation state corresponds to three successive proton transfers: proton export from MotA.E151 to the cytoplasm side, proton hopping from MotB.D22 to MotA.E151, and proton uptake from the periplasm side to MotB.D22. This corresponds to net single-proton transfer across the membrane. Notably, however, exports occur in a different position from hopping and uptake; specifically, for the first update from the cryo-EM structure, a proton is exported from MotA(chain A).E151 to cytoplasm, whereas proton hopping occurs in MotB(chain G).D22 to MotA(chain E).E151, followed by proton uptake by MotB(chain G).D22. Together, these three proton transfers resulted in a ∼36° rotation of the MotA pentamer. We address the plausible order of the three proton transfers in more detail later in this paper.

### Deciphering an optimal pathway

By varying the number of deprotonated sites in the MotA pentamer, we found that robust rotary motion occurred only when there were two deprotonated sites in the basal state. Specifically, we select the initial states [(C, E), ()]. However, there are several other possible explanations for the two deprotonated sites in the MotA pentamer. Here, we examined other cases.

The cases of the initial states [MotA, MotB] = [(C, E), (F)] are shown in Fig. S3a (this initial state does not satisfy the first propensity rule; however, the rule is matched after the first update). In this case, there should be proton uptake from periplasm to MotB(chain F).D22. Subsequently, a proton is transferred from MotB(chain G).D22 to MotA(chain E).E151 and proton export from MotA(chain A).E151 reaches the updated protonation state [(A, C), (G)]. The simulations for one turn with this setup resulted in robust rotary motions of approximately 36° per update, similar to the case of [(C, E), ()] (Fig. S3a).

Next, we examined another case in which the initial state [MotA, MotB] = [(B, E), (G)] (Fig. S3b). In this case, proton hopping occurred from MotB(chain F).D22 to MotA(chain B).E151, a proton exported from MotA(chain C).E151 to cytoplasm, followed by proton uptake from periplasm to MotB(chain G).D22 reaches the updated protonation state [(C, E), (F)]. In a simulation using this setup, we observed a robust rotary motion of ∼36° per update (Fig. S3b). We note, however, that in the initial protonation state [(B, E), (G)], the stable angle of the MotA pentamer in the simulation was ∼ -61° from the cryo-EM structure. Thus, although this type of update can induce the desired rotary motion of approximately 36° per update, the initial protonation state is less likely to be compatible with the cryo-EM structure.

Furthermore, we considered other cases, in which two MotA.E151 were deprotonated in the initial state: [MotA, MotB] = [(C, E), (G)], [(C, E), (F, G)], and [(D, E), (G)]. First, the scheme with the initial state [(C, E), (F, G)] does not satisfy the first propensity rule at any stage, and thus is unlikely (Fig. S3d). Secondly, for [(C, E), (G)] and [(D, E), (G)], the schemes require the uptake and export of two protons for a single update of the protonation state, which is unlikely. (Fig. S3ce).

Altogether, we found a robust rotary motion of the MotA pentamer with initial protonation states [MotA, MotB] = [(C, E), ()], [(C, E), (F)], and [(B, E), (G)]. Interestingly, these three states can be understood as different steps in the same pathway, making these states essentially synonymous. Specifically, [(C, E), (F)] is immediately followed by [(C, E), ()] for MotB(chain F).D22 receives a proton from periplasm. Moreover, the states [(B, E), (G)] are updated to reach [(C, E), (F)] (Fig. S3b) with a rotation of ∼ 36 °. Thus, although it is unclear which of the three states correspond to the cryo-EM structure at this stage, they all suggest the same pathway. Based on the satisfaction of all the three propensity rules and relatively small deviation of the rotary angle in the initial state, we suggest that, from our model analysis, the cyclic update of the protonation state from [MotA, MotB] = [(C, E), ()] is an optimal pathway.

### The free energy landscape supports the optimal pathway

To further understand the robust rotary motion identified in the optimal pathway, we analyzed the free-energy landscape of the state [MotA, MotB] = [(C, E), ()], as well as [(A, C, E), ()] as a control. We performed longer MD simulations for the two protonation states with eight equally spaced initial rotary angles and collected all the samples to calculate the equilibrated rotary angle distribution, from which we plotted the free energy landscape along the rotary angles in Fig. 3 (see the Methods section for more details). The landscape with [(C, E), ()] has one sharp basin near the cryo-EM structure at ∼ -22.5°, in addition to a well-separated basin at the pseudo twofold-symmetric angle at ∼ +147.5. With the symmetry of the 5:2 system, we expect that one-step update of the protonated state to [(A, C), ()] leads to the free energy basins at ∼ -22.5+36° and ∼ +147.5+36°. Because of the well-separated nature of the two basins, the MotA pentamer steadily rotated by 36° per update.

**Figure 3:**
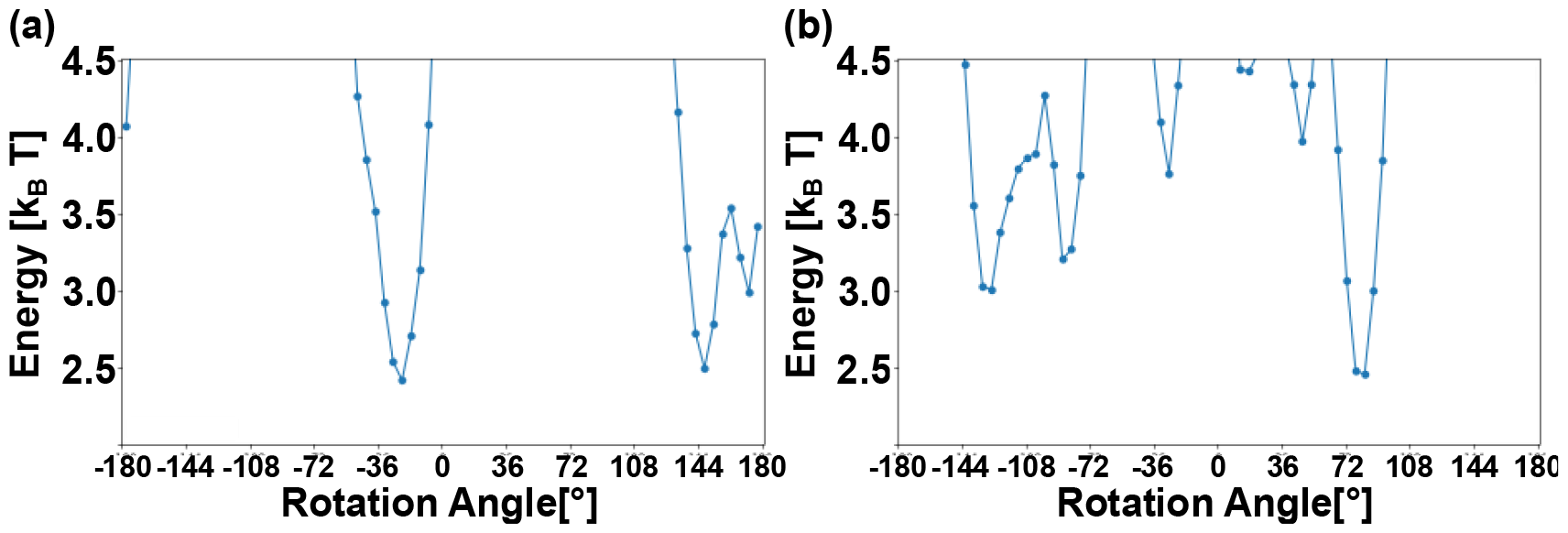
The free energy landscapes of the two protonation states. (a) The protonation state [(C, E), ()] having the two major basins. (b) The protonation state [(A, C, E), ()] having rugged and broader basins.

In contrast, the protonation state [(A, C, E), ()] possesses a broader and more rugged landscape, in addition to a deep basin of ∼ +82.5° (Fig. 3b), which is rather far from the cryo-EM structure. The following two points render this state less compatible with the cryo-EM structure: First, the deviation of the stable rotary angle from the cryo-EM structure was markedly large. In principle, if we assign the protonation state of [(E, B, D), ()], which corresponds to a two-step backward shift, we expect to obtain a deep free-energy basin near the cryo-EM structure. However, this mapping of the protonation state is not compatible with the proton uptake timing based on the cryo-EM structure and thus is unlikely. Second, we expect that the one-step update of the protonation state from [(A, C, E), ()] to [(A, B, D), ()] will lead to a similar free-energy landscape with a 36° shift to the right. Owing to the rugged nature of the landscape before and after the update, the transition from one to the next protonation state would not result in robust rotation of the MotA pentamer.

Thus, the underlying free-energy landscape supports robust rotary motions in the optimal pathway starting from [(C, E), ()] and irregular motions in other cases.

### Small translational motion of MotA in the rotation

The MotAB motor has the 5:2 symmetry of MotA and MotB, which is distinct from other well-known ion-driven rotary motors, such as the F_O_ motor in F-type ATP synthase. The latter contains only one a-subunit and species-dependent number of c-subunits thus having the n:1 symmetry. The 5:2 symmetry suggests that after a one-step update associated with a 36° rotation, the roles of the two MotB subunits must change. For example, the proton uptake by MotB(chain F).D22 leads to the first state shown in Fig. 2c, whereas proton uptake in the next step occurs in MotB(chain G). This alternating nature of the two MotB subunits can induce not only rotary motion, but also some translational motion along the MotB chain G/chain F axis (w-axis in Fig. 4a, center defined by the vector formed by two K15s of MotB). We plotted the translational motion along the w-axis for the two pathways, [MotA, MotB] = [(C, E), ()] and [(A, C, E), ()] (Fig. 4b-h) for the same trajectories used in Fig. 2h and i, respectively. In the [(C, E), ()] case, which showed the sustained rotation, a periodic pattern of the translational motion between w = +5 and -5 was observed (Fig. 4c). In contrast, in the [(A, C, E), ()] case, which did not exhibit a net rotation, this periodicity was disrupted (Fig. 4g).

**Fig. 4.**
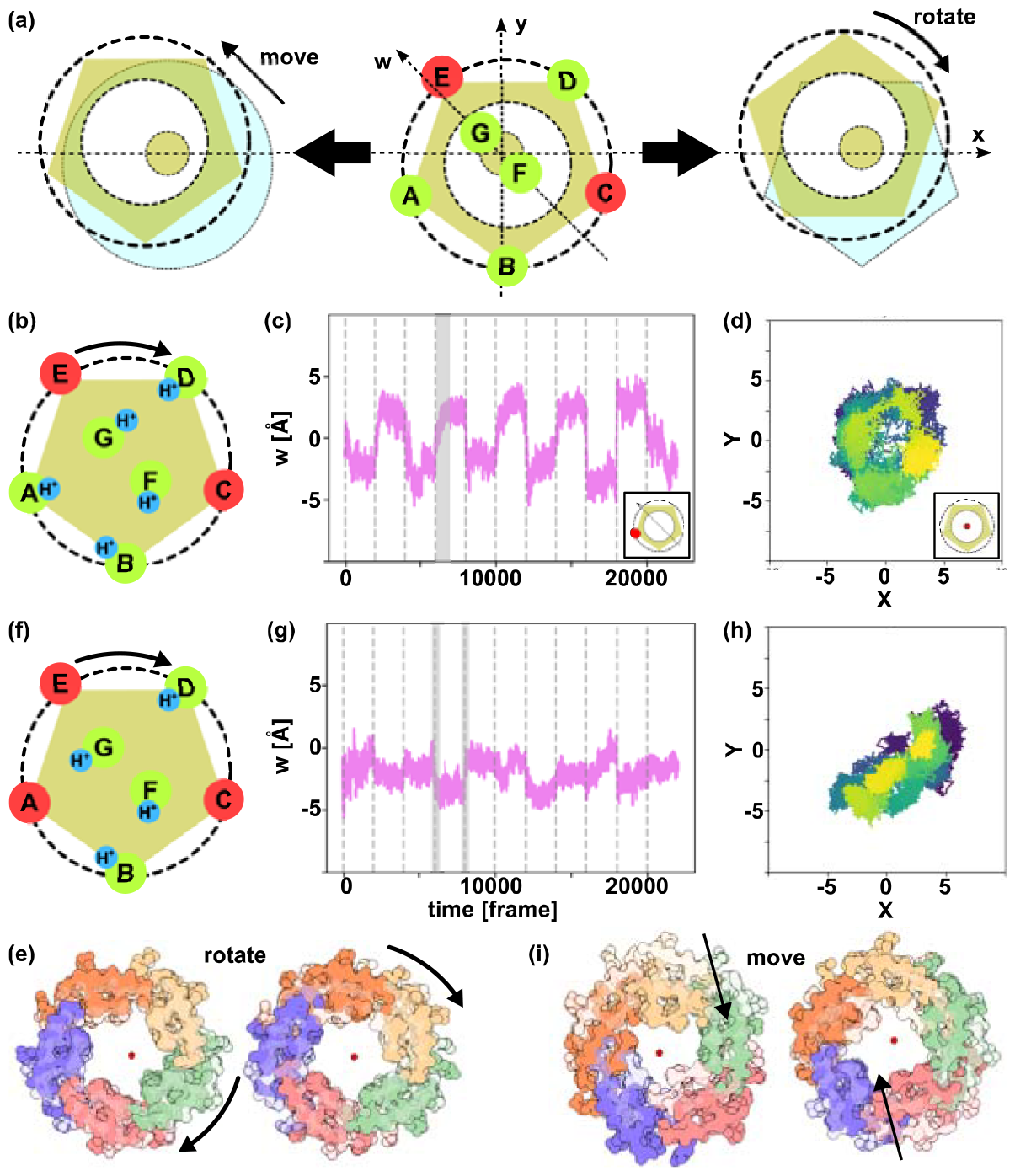
Trajectories of MotA’s center of mass. (a) Definition of the w-axis (center), with schematic representations of MotA exhibiting translational movement without rotation (left) and rotary movement (right). (b) Schematic diagram of [MotA, MotB] = [(C, E), ()], corresponding to Fig. 2c. (c) Trajectory of MotA(chain A).E151 along the w-axis for [(C, E), ()], with the structure in the gray area shown in (e). (d) Trajectory of the center of mass coordinates for E151 of Chains A-E in the trajectory from (c). (e) Snapshots of MotA corresponding to the gray area in (c); both semi-transparent snapshots are at Frame 6200, with the left opaque at Frame 6400 and the right at 6800. (f) Schematic diagram of [MotA, MotB] = [(A, C, E), ()], identical to Fig. 2d. (g) Trajectory of MotA(chain A).E151 along the w-axis for [(A, C, E), ()], with the structure in the gray area shown in (i). (h) Trajectory of the center of mass coordinates for E151 of Chains A-E in the trajectory from (g). (i) Snapshots of MotA corresponding to the gray area in (g); the left semi-transparent is at 5900 frames, opaque at 6100 frames, while the right semi-transparent is at 7900 frames, opaque at 8100 frames.

Additionally, to directly follow the positional changes in MotA relative to MotB, the center of mass of the five E151s of the MotA pentamer was calculated, and their trajectories were plotted on the xy-plane. Although Fig. 4d shows the center of mass of MotA forming a neat donut shape, Fig. 4h shows a disordered plot (all plots are shown in Fig. S4).

These results suggest that rotary motion is associated with the small seesaw-like translational motion of the MotA pentamer relative to the MotB dimer.

### Analysis of MotAB mutants

Our modeling assumes the three residues that play major roles, MotA.E151, MotB.D22, and MotB.K15. Here, we examined effects of mutations in these residues. Both MotA.E151 and MotB.D22 are the proton-carrying sites, while MotB.K15 is a key residue to generate a rotational torque. We performed MD simulations for the four mutant constructs: MotA(chain D).E151A, MotB(chain F).D22A, MotB(chain F).K15A, and MotB(chain F and G).K15A. `Note that mutations are included only in the specified subunits.`All the simulations used the same setting as in the [(C, E), ()].

First, the mutant MotA(chain D).E151A did not rotate as the WT (Fig. S5a). Of the 10 updates, we observed ∼36° rotation in some limited cases. Notably, we introduced the mutation E151 only in one MotA subunit (chain D). In some rotary steps, MotA(chain D) is not involved in the proton transfer at all. Thus, it was expected that these steps were not affected by this mutation. The results are consistent with this expectation. Yet, the overall rotation is rather severely affected by this single mutation. Note, however, that the E151 moiety, in reality, contains E151 and the backbone of G150. Thus, the mutant E151A in real molecules may have different effects from the current results.

Second, the mutant MotB(chain F).D22A did not rotate at all (Fig. S5b). In this case, MotB.D22A does not receive any proton from periplasm, inhibiting its rotation. Even if only one of the two MotB chains have this mutant, the overall rotation is completely inhibited. In previous experiments, the corresponding mutation (D33A) for *S. enterica* MotAB showed the loss of function (37,38). Our mutant simulation is consistent with this result.

Third, the mutant MotB(chain F).K15A, interestingly, exhibited partial rotation, albeit unstable (Fig. S5c). This is probably due to the fact that the proton transfer itself is not inhibited by this mutation. The source of rotary torque was only halved, but not disappeared.

Finally, the mutant MotB(chains F and G).K15A no longer rotated (Fig. S5d). While proton transfer itself was not inhibited, the rotational torque was not generated at all. Notably, MotB molecules have disordered N-terminal tails which contain some positively charged residues. If K15 is mutated, some N-terminal disordered region may have similar roles. Our current modeling did not contain these disordered regions

### MotB to MotA proton hopping before proton export from MotA to cytoplasm

Our model simulation suggests that the pathway starting from the protonation state [(C, E), ()] is most compatible with the cryo-EM structure and robust rotation. In the simulations so far, we updated the protonation state from [(C, E), ()] to [(A, C), ()], resulting in a 36° rotation. However, this update contains three proton movements: (1) Proton hopping from MotB(chain G).D22 to MotA(chain E). E151; (2) proton export from MotA(chain A).E151 to the cytoplasm, and (3) proton uptake from the periplasm to MotB(chain G).D22. What is the order of the three-proton movements? Which of the three proton motions is coupled to rotation? MotB(chain G).D22 cannot accept two protons simultaneously and thus the movement (3) must occur after process (1). Moreover, process (3) is expected to occur after MotA pentamer rotation because of the side-chain reorientation of MotA.F186 is essential for proton uptake in process (3). Thus, Movement (3) is most likely the last step of the three movements. However, it is unclear whether (1) or (2) occurred first (Fig. 5a,b).

**Fig. 5:**
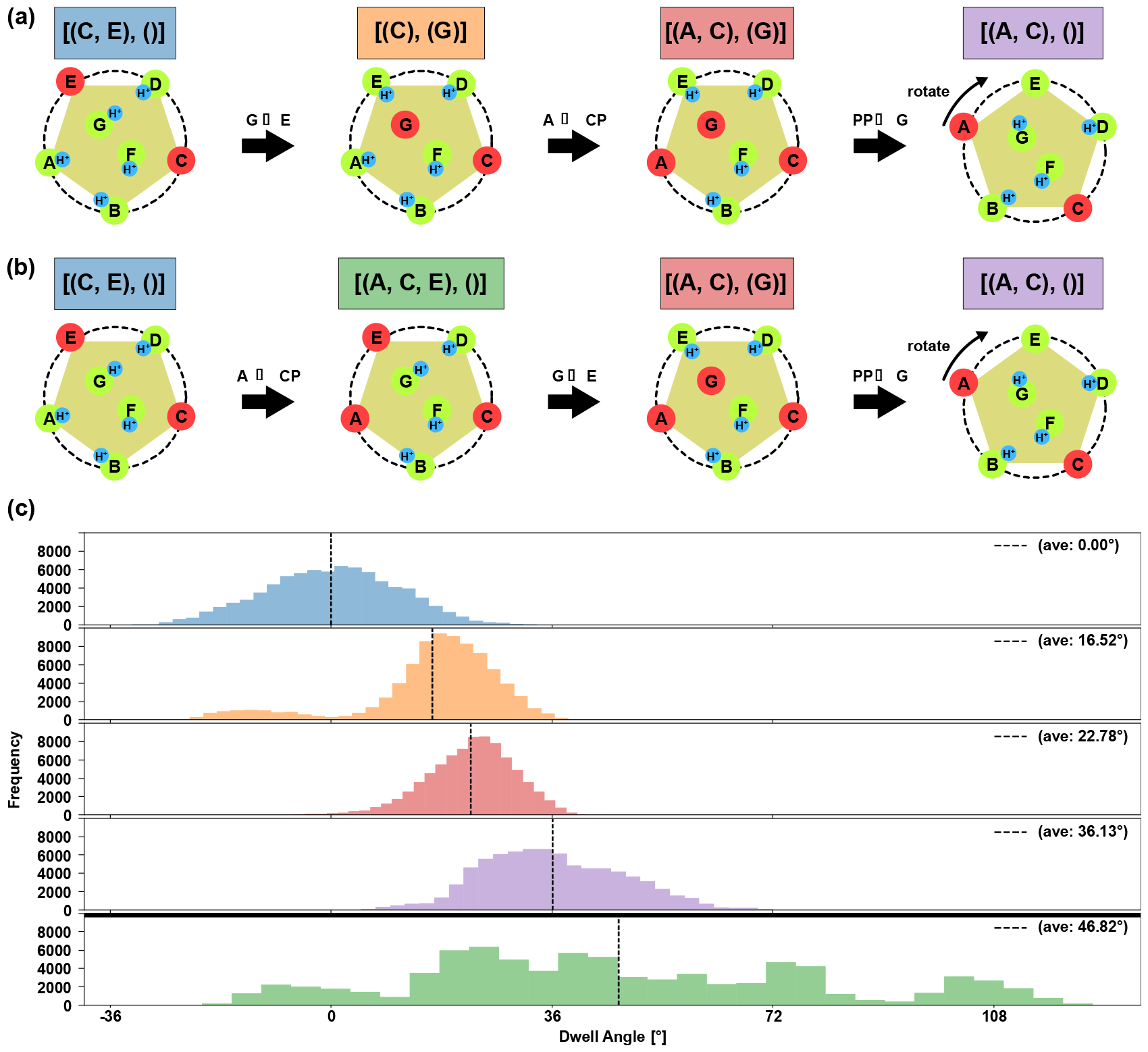
Proton transfer coupled rotary pathways of MotAB. Two alternative pathways (a) and (b) are considered to decipher the proton transfer pathways. (a) Proton transfer occurs in the order of the proton hopping -> the proton export -> the proton uptake. (b) Proton transfer occurs in the order of the proton export -> the proton hopping -> the proton uptake. (c) The rotary angle distributions for each protonation state. The color of each plot corresponds to that in the two pathways in (a)(b). The angle is plotted as the difference from the mean position of the initial state [(C, E), ()]. The average angles for each color are listed at the right.

To this end, we addressed the order of the first two proton movements: (1) and (2). If proton hopping (1) is followed by proton export (2), `then the pathway is from` [(C, E), ()] to [(C), (G)], [(A, C), (G)], and [(A, C), ()] (Fig.5a). Otherwise, the pathway was `from` [(C, E), ()] to [(A, C,E), ()], [(A, C), (G)], [(A, C), ()] (Fig. 5b). To infer the rotary motion along the two pathways, we calculated the rotary angle distributions in the five states: [MotA, MotB] = [(C, E), ()], [(C), (G)], [(A, C, E), ()], [(A, C), (G)], and [(A, C), ()]. Each protonation state was set at the initial configuration stage, and simulations were performed five-fold the MD time steps used previously while maintaining a single protonation state. The rotary-angle distributions for the five states are shown in Fig. 5c, where the change in the rotary angle was calculated relative to the average dwell angle of [(C, E), ()].

The results clearly show stepwise rotations following the first scheme from [(C, E), ()] to [(C), (G)] to [(A, C), (G)], and [(A, C), ()]. The first proton hopping shifts the mean rotary angle by 16.5°, followed by proton export, which further shifts the mean angle by 6.2°. The final step of proton uptake resulted in a 13.4 °shift in the mean angle. The results suggest that each proton movement partially contributes to the rotary motion, making the rotation smooth and steady.

However, if the second scheme is used, the second stage has a protonation state [(A, C, E), ()], of which the free energy landscape was discussed above. [(A, C, E), ()] have a wider and less organized distribution of rotary angles, making the second scheme less likely to obtain robust rotation.

Therefore, we conclude that a smooth rotary motion occurs along the pathway shown in Fig. 5a, where protons hop from MotB(chain G).D22 to MotA(chain E).E151 occurs before the proton export from MotA(chain A).E151 to cytoplasm, followed by proton uptake from periplasm to MotB(chain G).D22.

## Discussion and Conclusion

We built a protonation-dependent computation model of MotAB based on the structure of MotAB from the gram-negative *Campylobacter jejuni* (PDB ID: 6YKP) and conducted protonation-dependent MD simulations to investigate the rotary mechanisms and the proton transfer coupled rotation pathway. Upon comprehensive examination of numerous protonation states, we found that a robust ∼36° rotary motion per proton transfer across the membrane occurred for the pathway starting from [(C, E), ()], with two of the five MotA.E151 deprotonated in the basal state. By deciphering the individual proton transfers, we suggested the most plausible proton transfer coupled pathway, as shown in Fig. 5a.

It would be interesting to compare the rotary mechanism of the two ion-driven rotary motors, the MotAB motor and the F_O_ motor of F-type ATP synthase. The F_O_ motor is made of an a-subunit serving as a stator and a ring-shaped c-subunit oligomer with varying number of c-subunits, which serves as a rotor. The c-subunit contains the highly conserved acidic residue, glutamate or aspartate, in the middle of its transmembrane helix facing to the interface between the stator and the rotor. Two half-channels are found near the interface between the a-subunit and the c-ring, each connecting the c-subunit acidic residue to either cytoplasm or periplasm (for the bacterial one) side. This combination of two half-channels and one well-conserved acidic residue in the middle of the transmembrane helix facing towards the interface seems to be shared by the MotAB motor. A noticeable difference between the two systems is their stoichiometry. The F_O_ motor has an n:1 stoichiometry, whereas MotAB has a 5:2 stoichiometry. Because the a-subunit in F_O_ is a monomer, all relevant proton transfers occur at the interface between the a-subunit and the c-subunits facing the a-subunit. Thus, the reaction is localized to the single a-subunit neighbor. In contrast, the 5:2 MotAB configuration resulted in more delocalized reactions. In the pathway that we suggest as plausible (Fig. 2c), the first 36° rotation is associated with proton transfer near the MotB(chain G) side, whereas the second 36° rotation is due to the oppositely facing MotB(chain F). The first rotation involved proton motion in chains A, E, and G, whereas the second rotation was mediated by chains C, D, and F, which were mutually exclusive. This alternate use of subunits in successive rotations is a key feature of 5:2 stoichiometry.

One major limitation of the current modeling is the proton (or hydronium ion)-carrying site next to the well-conserved MotB.D22. For simplicity, we designated it MotA.E151, although a similar role can be played by the backbone carbonyl of MotA.G150, as previously mentioned. Notably, site 151 is modestly conserved (See Figure S1 of (5)). Both glutamic and aspartic acids are present in many, but not all, related species. MotA.G150 was more conserved. More importantly, MotA.P154 is strictly conserved, suggesting that the backbone carbonyl group at site 150 does not form a hydrogen bond in the alpha helix, thus favoring a kinked structure. This moiety can accommodate hydronium ions that serve as proton-carrying sites. Obviously, a more detailed analysis with full atomic resolution is required.

As a first step in the computational modeling of MotAB, we employed a relatively simple model; thus, there are some technical limitations to be resolved in the future. First, the simulations are based on a coarse-grained model. Although the parameters of the coarse-grained AICG2+ model were derived from atomistic interaction energies, the model did not explicitly represent the side chain. Specifically, with atomistic modeling, the side-chain reorientation of MotA.F186, as well as those of glutamic and aspartic acids, can be addressed. This side-chain repositioning is expected to facilitate proton transfer. Secondly, the simulation did not explicitly include lipids or water molecules. In the current approach, we minimally modeled the membrane environment by restraining the MotA pentamer to a planar geometry. However, the rotation of the pentameric molecule induces remodeling of the lipid membrane, which could affect rotary dynamics. Solvation and water dynamics play major roles defining the half-channels and proton/hydronium ion-carrying sites. All-atom MD simulations with explicit lipids and water molecules are highly desirable. Finally, in the proposed approach, the timing of updating the protonation states does not depend on the structural state. In the main simulations, we updated the protonation state every 2000 frames, regardless of the rotary state. However, proton uptake from PP to MotB is crucially dependent on the reorientation of F186, which is expected to depend on the rotary angle of the MotA pentamer. Therefore, it is highly desirable to include a dynamic proton-hopping process, which depends on protein configuration, instead of updates at predetermined timings. Previously, for the Fo motor in F-type ATP synthase, we employed a hybrid Monte Carlo(MC)/MD simulation to simulate the proton-transfer-coupled rotary motion of the c-ring (34). In future, we hope to perform similar hybrid MC/MD simulations for MotAB.

Despite these challenges, the current study is the first to elucidate that robust rotary motion can be achieved by updating the two deprotonated sites in the MotA pentamer. Additionally, insights into the rotary pathways of MotAB from other species not covered in this study have been obtained.

## Materials & Methods

### Model building

We used the MotAB structure of gram-negative *Campylobacter jejuni* (PDB ID: 6YKP) (5) as the reference structure for the MD simulations. The simulation system contain residues 1-255 of MotA chains A–E (UniProtKB: A0A0H3PAV1) and residues 15-39 of MotB chains F and G (UniProtKB: A0A0H3PBX6)(Fig. 1bc). In this analysis, we used the following coordinate system: First, we defined the geometric center of the N-terminal residues of MotB chains F and G. Similarly, we defined the geometric center of the C-terminal residues. The latter was set as the origin. The vector from the C-terminal center to the N-terminal center was set to the z-axis. We then set the x-axis such that MotA(chain C).A185 was on the x-axis.

### MD simulation setting

The MD simulations used a residue-resolution coarse-grained model with a single particle placing at the C_α_ position of each amino acid. We used the force field AICG2+ (40,41), which has long been used for a broad range of large-scale protein simulations because of its high speed and reasonable accuracy (34,42,43). The parameters therein were set based on fully atomistic interactions in the reference structure to reproduce the protein dynamics around the reference structure. The electrostatic interactions between MotA and MotB played a key role in this simulation. Therefore, we used the RESPAC method to locate partial and optimal charges on every surface amino acid so that the electrostatic potential around the protein surface by coarse-grained residues reproduced that of a fully atomistic model (44). Notably, RESPAC charge depends on the protonation state. For each protonation state, we created a structural model and obtained the RESPAC charges. The RESPAC charges were calculated for chain C of MotA and chain G of MotB, and were used for all the MotA and MotB subunits, respectively. Although MotAB is a membrane protein, we did not explicitly include lipid molecules. Instead, we fixed the positions of the N- and C-terminal residues of MotB chains F and G to serve as axles and restrained the MotA transmembrane region by the following addition potentials. We first identified that MotA.I21 and MotA.K64 are located near the boundary of the membrane region and just inside and outside the membrane, respectively. We applied a differentiable penalty function that takes large positive values when MotA.I21 is outside the membrane plane or when MotA.K64 is inside the membrane plane. Thus, the position of the MotB dimer is fixed, whereas the MotA pentamer can freely rotate within an effective two-dimensional space.

In this study, we updated the states of seven protonatable residues in MotAB, two D22 residues in MotB chains F and G, and five E151 residues in MotA chains A-E. Although these residues are negatively charged in their deprotonated states, their charges are zero in the protonated state. In all simulations for one turn, we updated the protonation state ten times, and for each protonation state, 2000×10^4^ MD steps were simulated. Thus, the overall simulation time was 11×2000×10^4^ MD steps. For convenience, we treated 1×10^4^ MD steps as 1 frame. We repeated the simulation of the same setup ten times with different random numbers. For Fig.3, we conducted 10^8^ MD step simulations with the fixed protonation state 10 times starting from 45n° rotated configurations with the integer 0 ≤ n ≤ 8. The sampled configurations were collected to obtain the free energy landscape. As shown in Fig. 5c, we conducted a 10^8^ MD step simulation with a fixed protonation state.

We used CafeMol version 2.1 for the simulations (39). We employed Langevin dynamics at 323 K and set the friction coefficient to 2.0 (in CafeMol units). Default settings were used for all the others.

### Analysis

The rotation angle of the MotA pentamer was determined by aligning the pentagons formed by the E151 residues of chains A-E. First, the center of mass of this pentagon was calculated in all steps, and the molecules were translocated along the xy-plane such that the center of mass matched that of the initial structure. The structure was then rotated around the z axis to minimize the root-mean-square deviation of the five E151 residues from those of the initial structure. The rotation angle was set to that of the MotA pentamer.

## Data Availability

The cryo-EM structures used in this study were downloaded from the Protein Data Bank under PDB ID 6YKP. All MD simulations in this study were performed using the CafeMol software. The data were downloaded from https://www.cafemol.org.

## Author Contribution

S.K. conceived and designed the project; S.K. developed the simulation code, performed the simulations, analyzed the data, and assembled the figures; S.K. and S.T. discussed the results and all authors were involved in the manuscript writing process.

## Acknowledgements

We would like to thank Toru Niina at RIKEN BDR for their support throughout this collaboration. We also thank Tomoko Furuya for their secretarial assistance.

