## Supplementary material for "Theoretical insights into rotary mechanism of MotAB in the bacterial flagellar motor": SI Figures

### Supporting Information

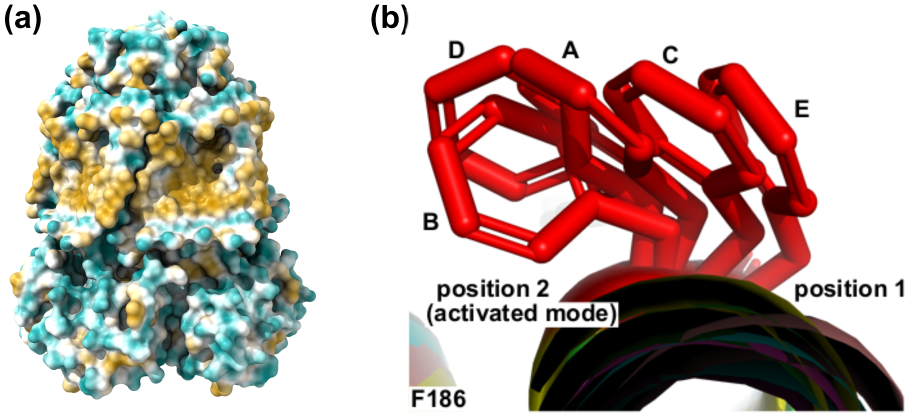


**Fig. S1 Structure.** (a) Hydrophobic interaction surface. (b) Orientation variations of F186.

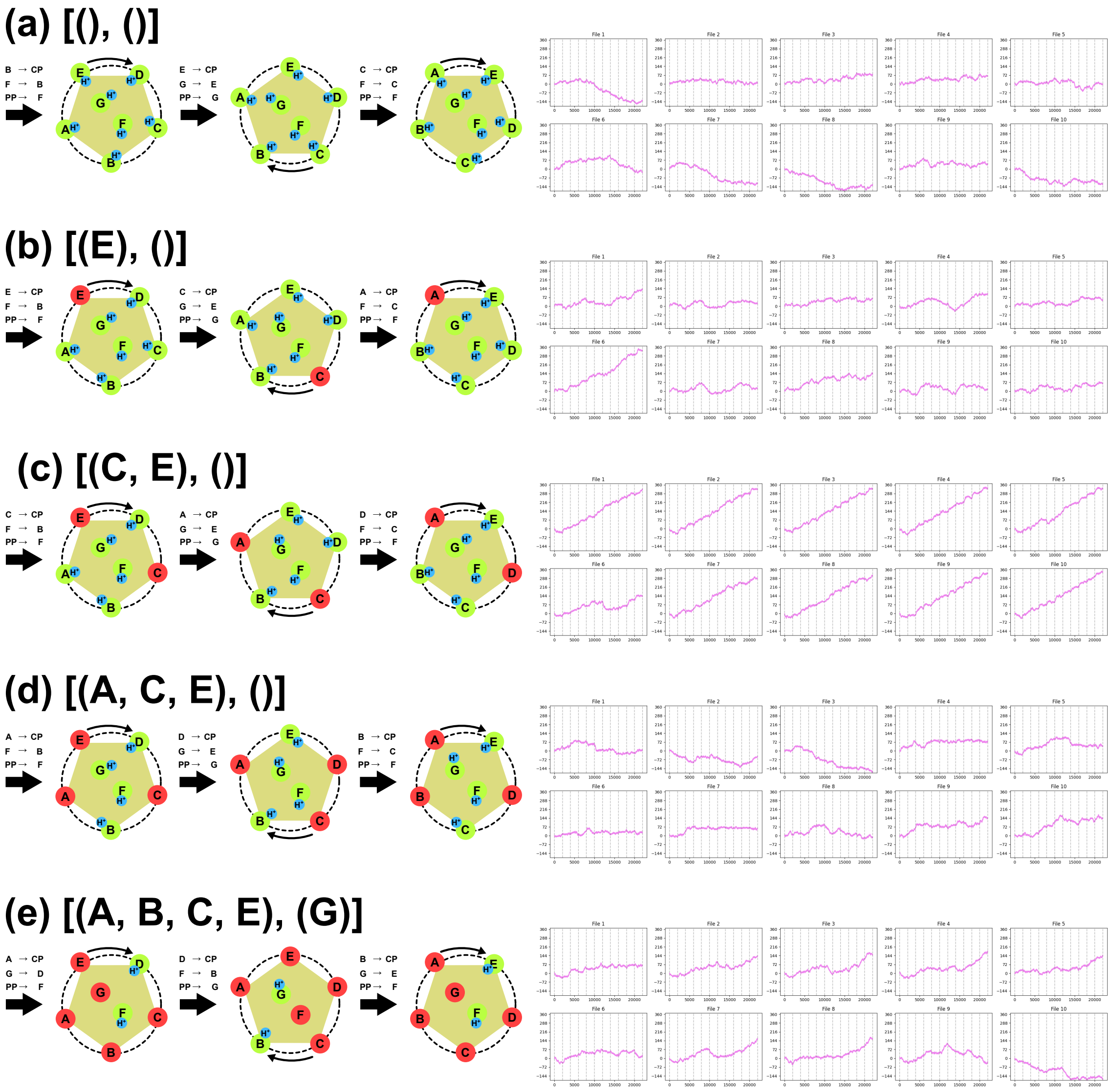


**Fig. S2 Entire trajectory of the pathway in Figure. 2.** Representative pathway diagrams and corresponding entire trajectories for MotA with 0, 1, 2, 3, and 4 deprotonated chain numbers. (a) [MotA, MotB] = [(), ()]. (b) [(E), ()]. (c) [(C, E), ()]. (d) [(A, C, E), ()]. (e) [(A, B, C, E), (G)].

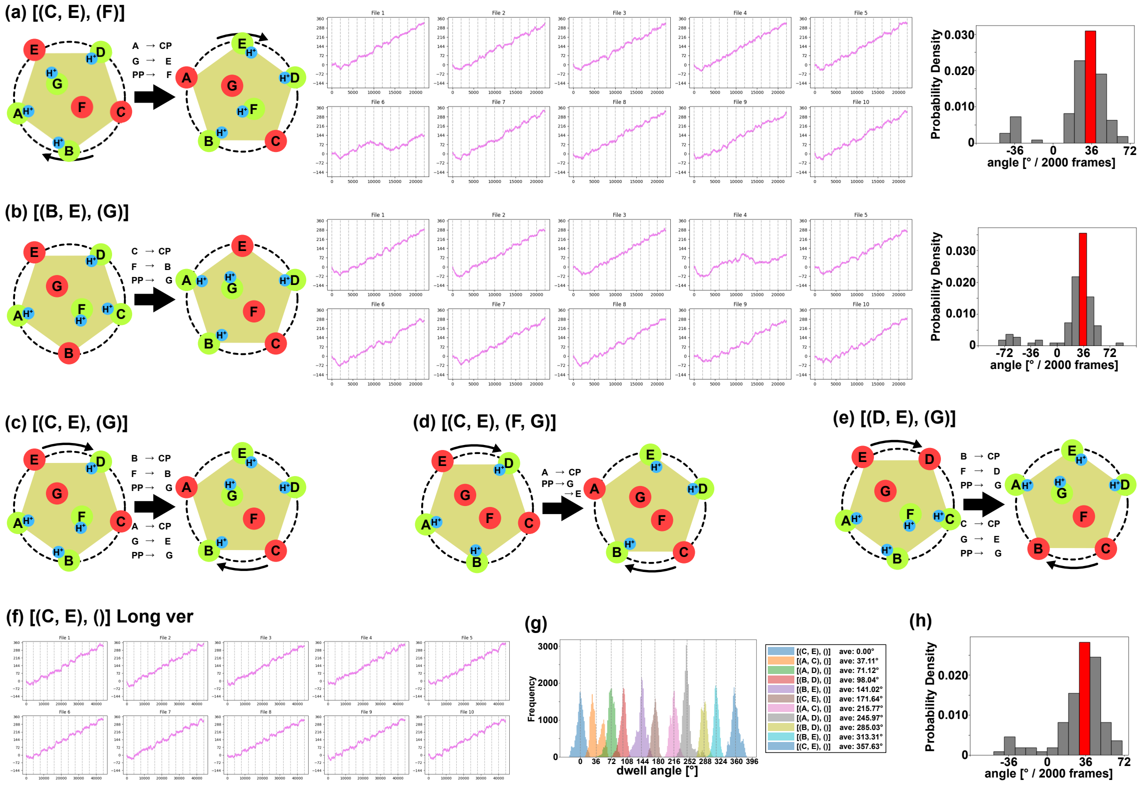


**Fig. S3: Alternative Pathways with Two Deprotonated Chains in MotA.** The pathway schematics. Red and green indicate deprotonation and protonation states, respectively, and the blue indicates proton. (a) [MotA, MotB] = [(C, E), (F)]. Left; pathway schematics. Middle; Entire rotational trajectory. Dotted lines indicate every 2000 frames. Right; Histograms of the rotation angle for every 2000 frames. The 36-degree rotation bin is highlighted in red. (b) [MotA, MotB] = [(B, E), (G)]. The pathway schematics, trajectory, and histogram. (c) [(C, E), (G)] pathway. (d) [(C, E), (F, G)] pathway. (e) [(D, E), (G)] pathway. (f) Entire rotational trajectory under 4000 frames per 1 state version of [(C, E), ()]. Dotted lines indicate every 4000 frames. (g) The rotary angle distributions for each protonation state. The color of each plot corresponds to the MotA protonation state. The average angles for each color are listed at the right. (h) Histogram of the rotary angle for every 4000 frames.

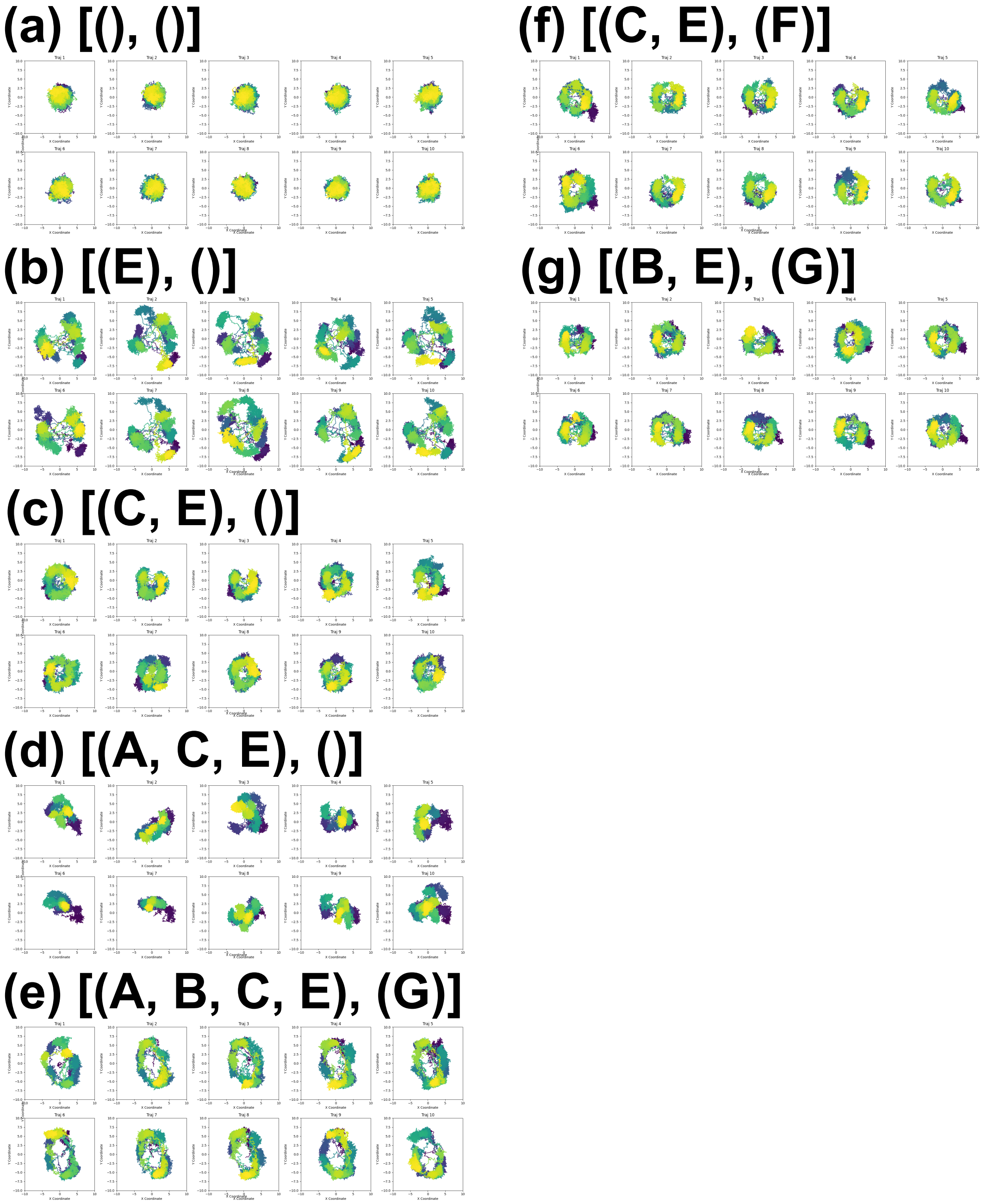


**Fig. S4 Trajectories of MotA center-of-mass for all paths.** (a) [MotA, MotB] = [(), ()] (b) [(E), ()] (c) [(C, E), ()] (d) [(A, C, E), ()] (e) [(A, B, C, E), (G)] (f) [(C, E), (F)] (g) [(B, E), (G)]

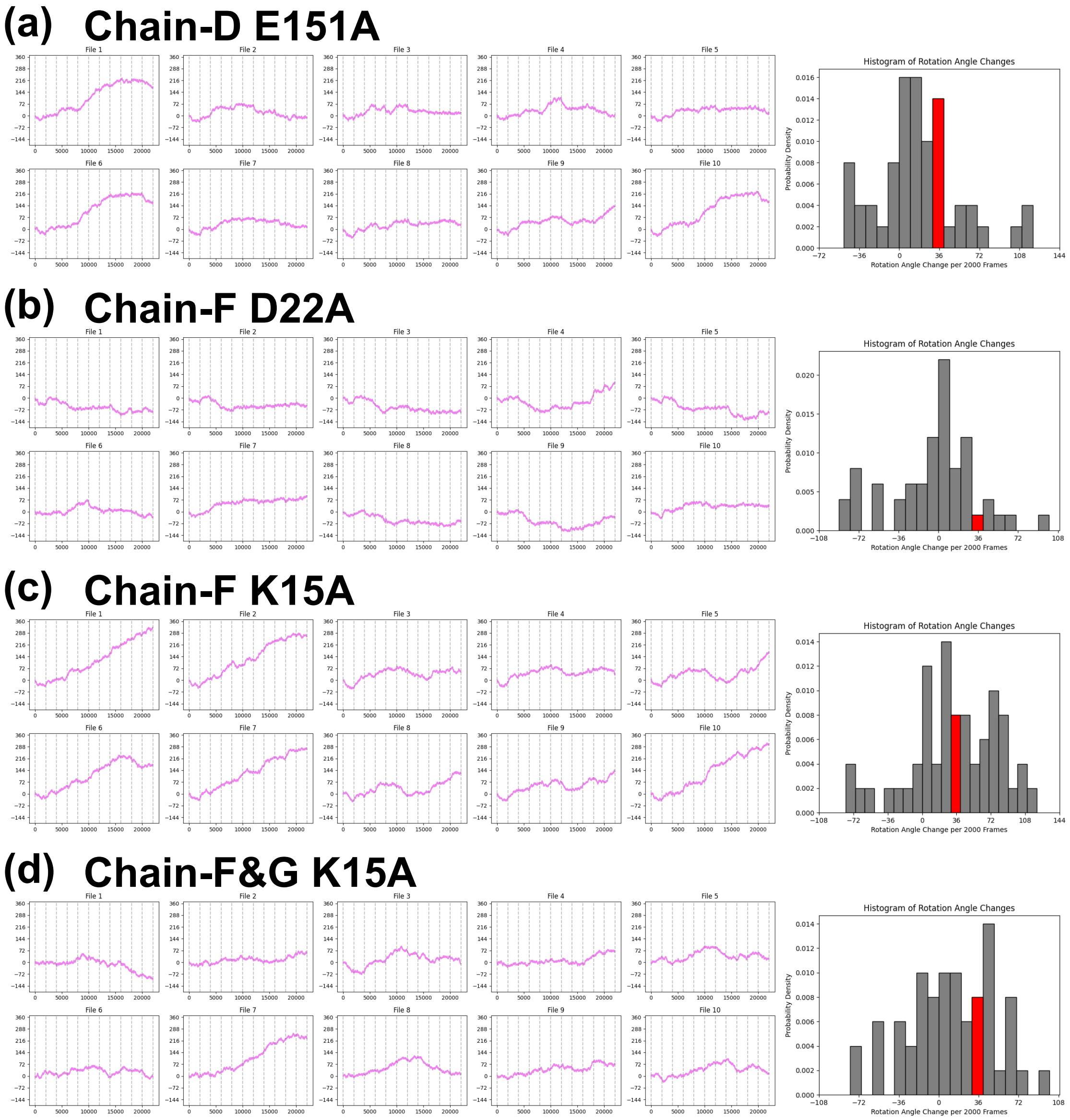


**Fig. S5: Simulations of mutant MotAB rotations in the [(C, E), ()] setting up.** Left: Entire rotary angle trajectories. Dotted lines indicate every 2000 frames. Right; Histograms of the rotation angle for every 2000 frames. The 36° rotation bin is highlighted in red. (a) MotA(chain D).E151A (b) MotB(chain F).D22A (c) MotB(chain F).K15A (d) MotB(chain F and G).K15A.

**Movie S1: Typical movie of [(C, E), ()].** Colour of each Chain is the same as in Fig. 1c. Clockwise rotation is forward; E151 in MotA and D22 in MotB and K15 in MotB are shown as spheres; K15 is always blue; E151 in MotA and D22 in MotB are green for protonated states and red for deprotonated states.

**Movie S2: Typical animation of [(A, C, E), ()].** The drawing method is the same as in Movie S1.
